## Supplemental Figures 1-29 for "A TOPBP1 Allele Causing Male Infertility Uncouples XY Silencing Dynamics From Sex Body Formation"

### Supplemental Figure 1

***Topbp1***$^{+/+}$

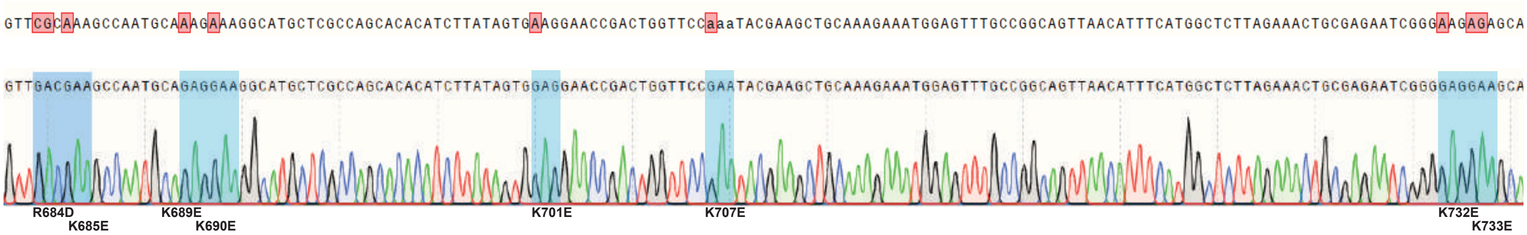


***Topbp1***$^{B5/B5}$

Supplemental Figure 2

A

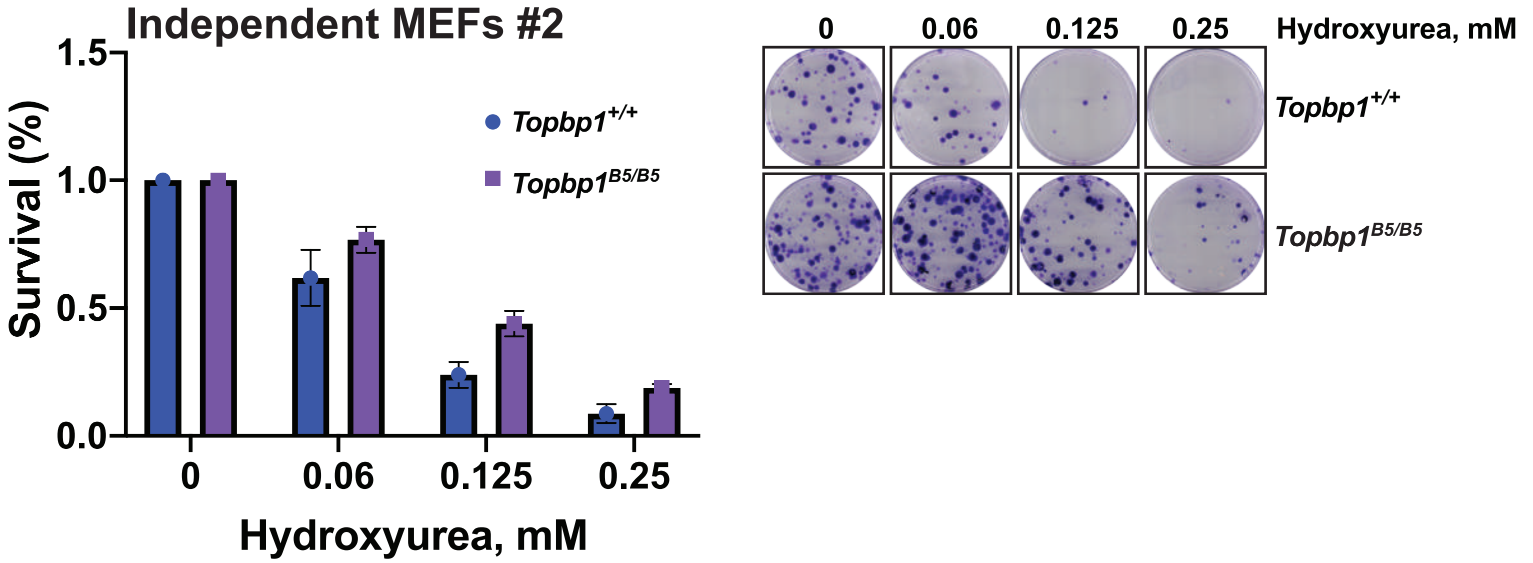


B

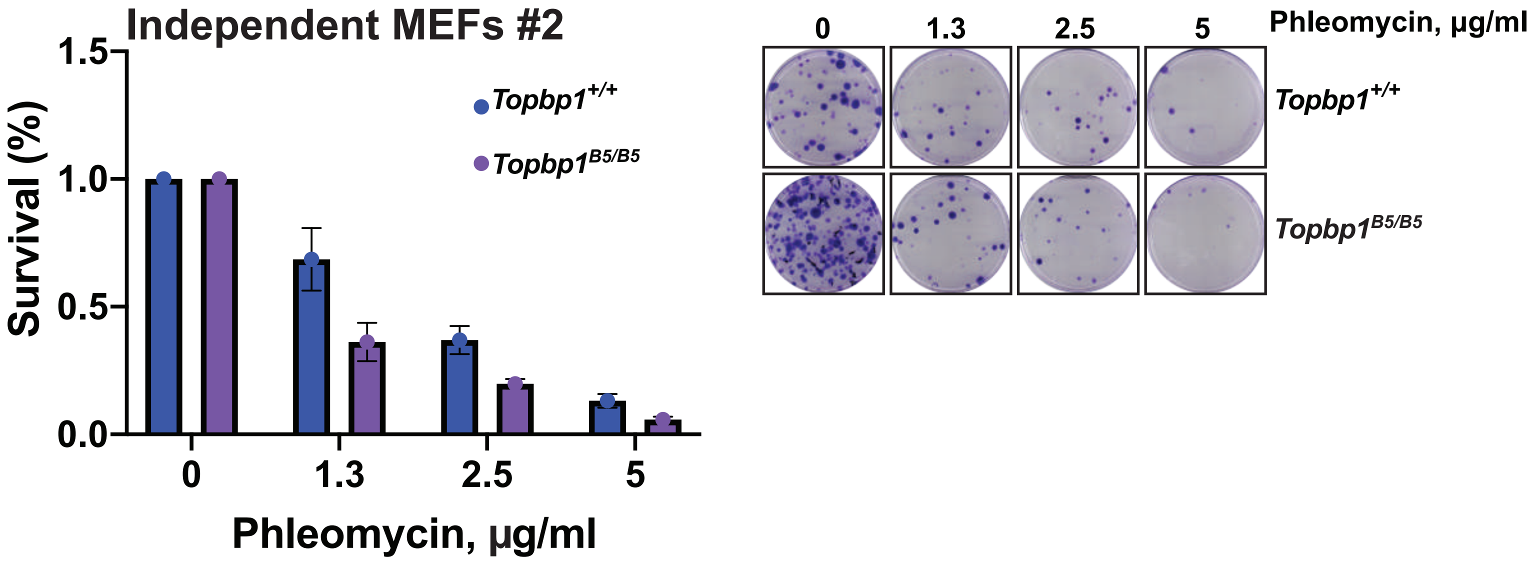

#### Supplemental Figure 3

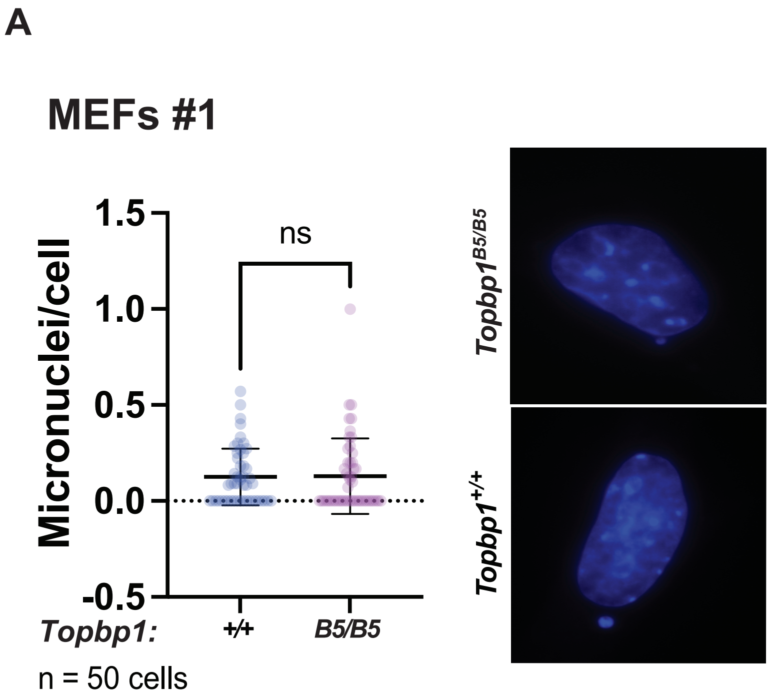


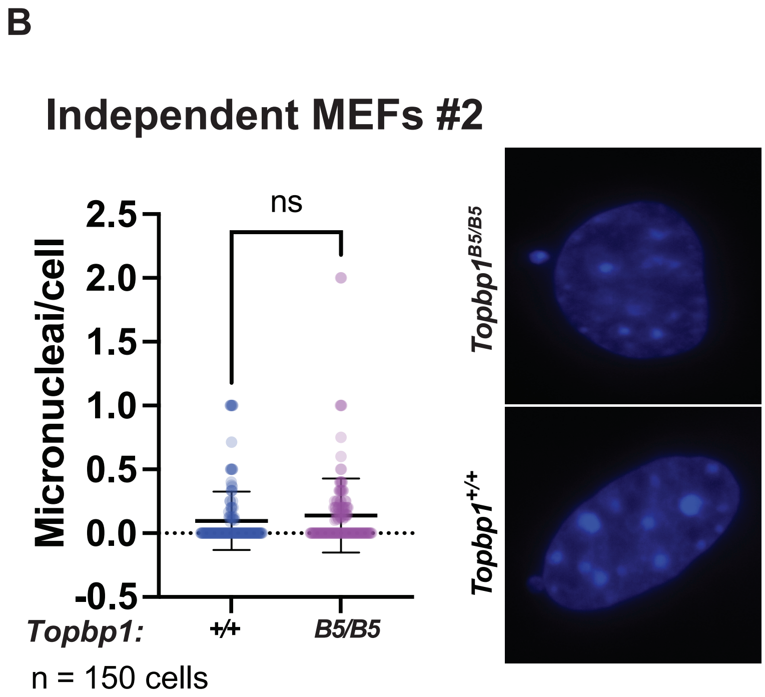

### Supplemental Figure 4

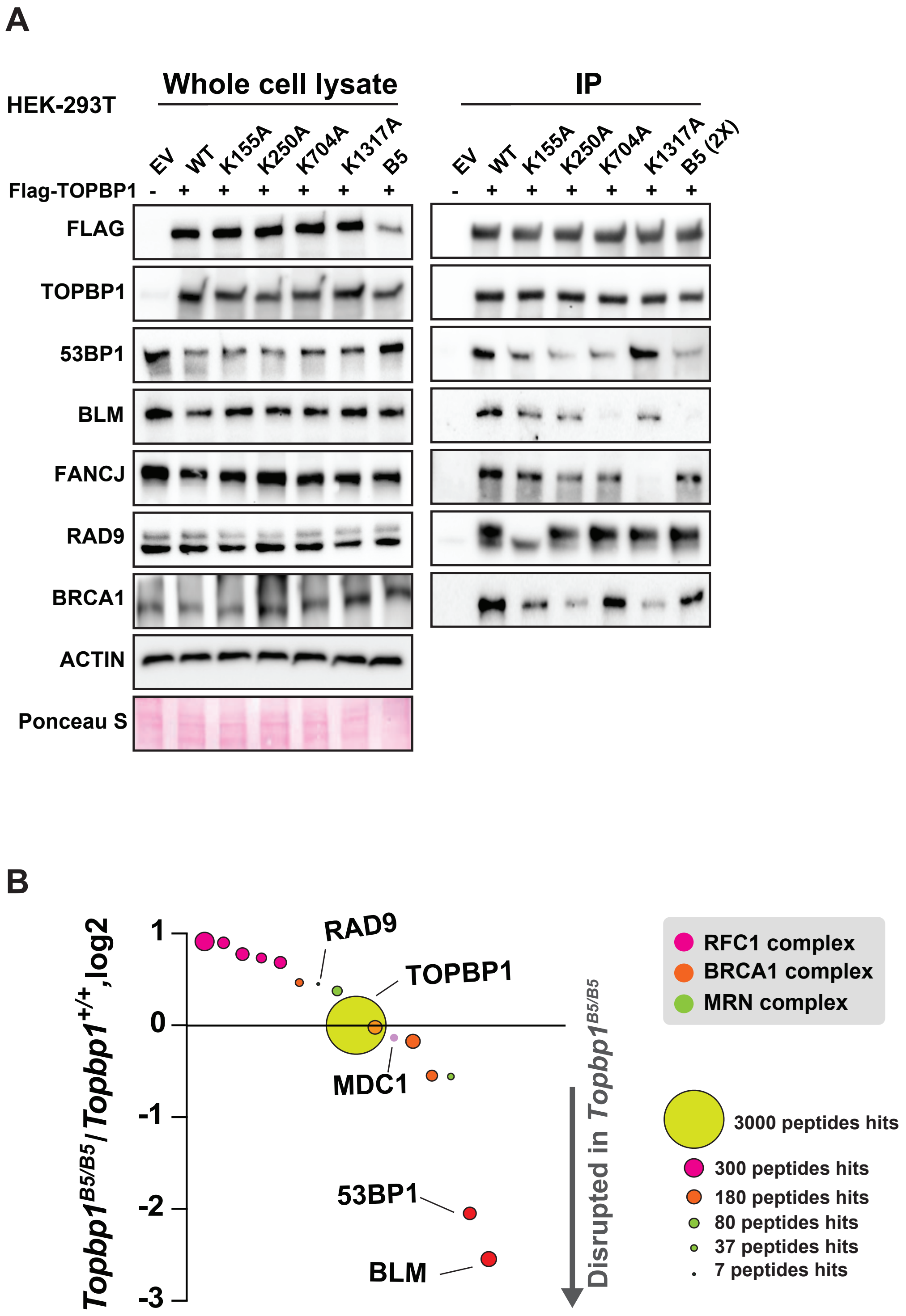

Supplemental Figure 5

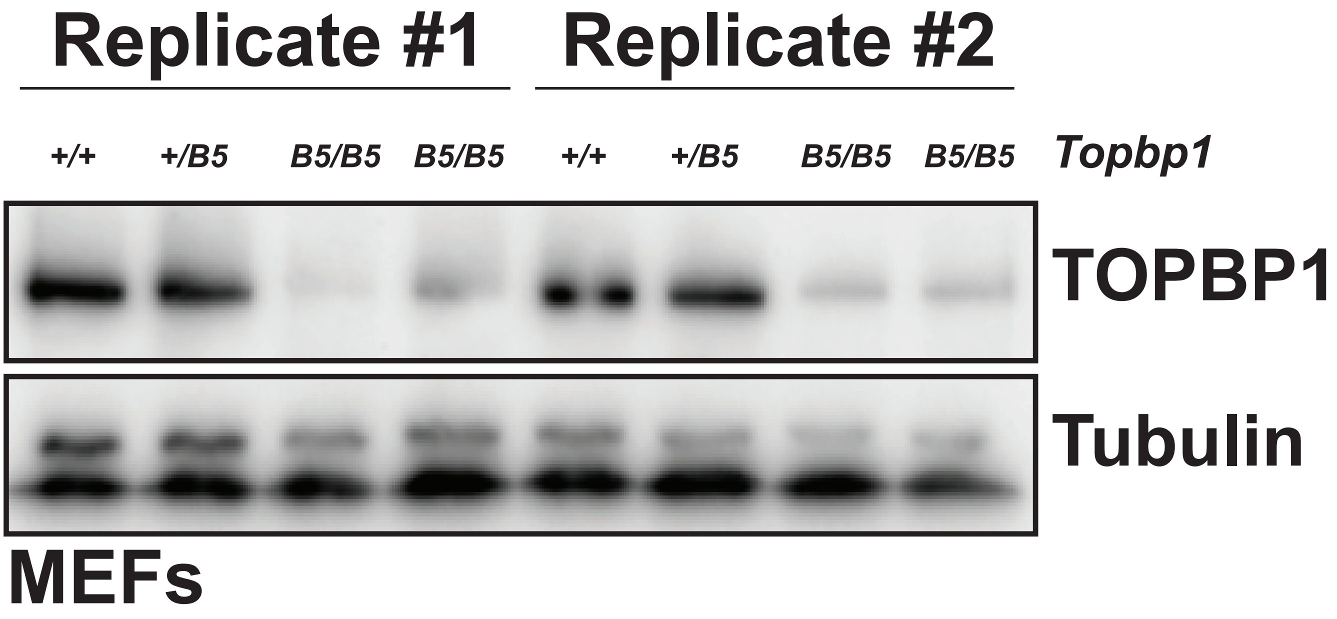

### Supplemental Figure 6

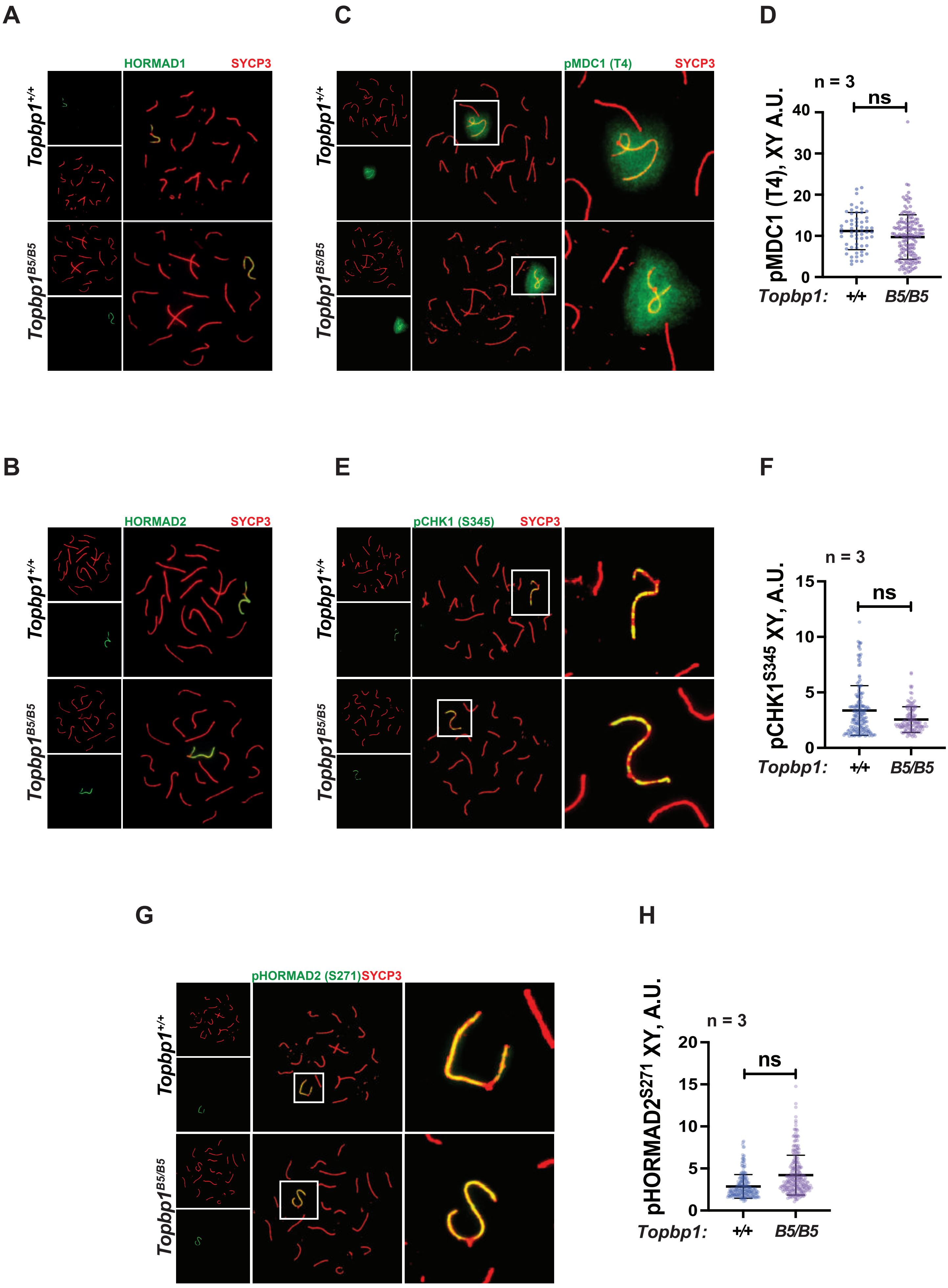

**Supplemental Figure 7**

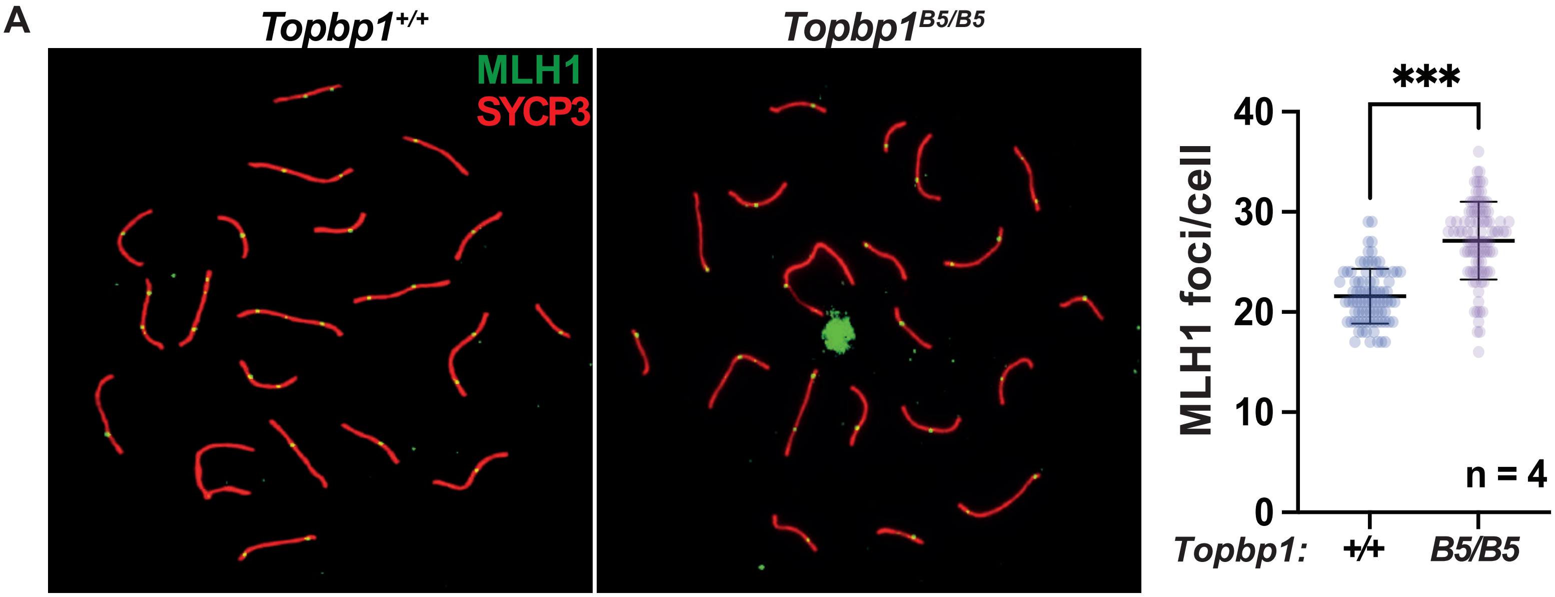


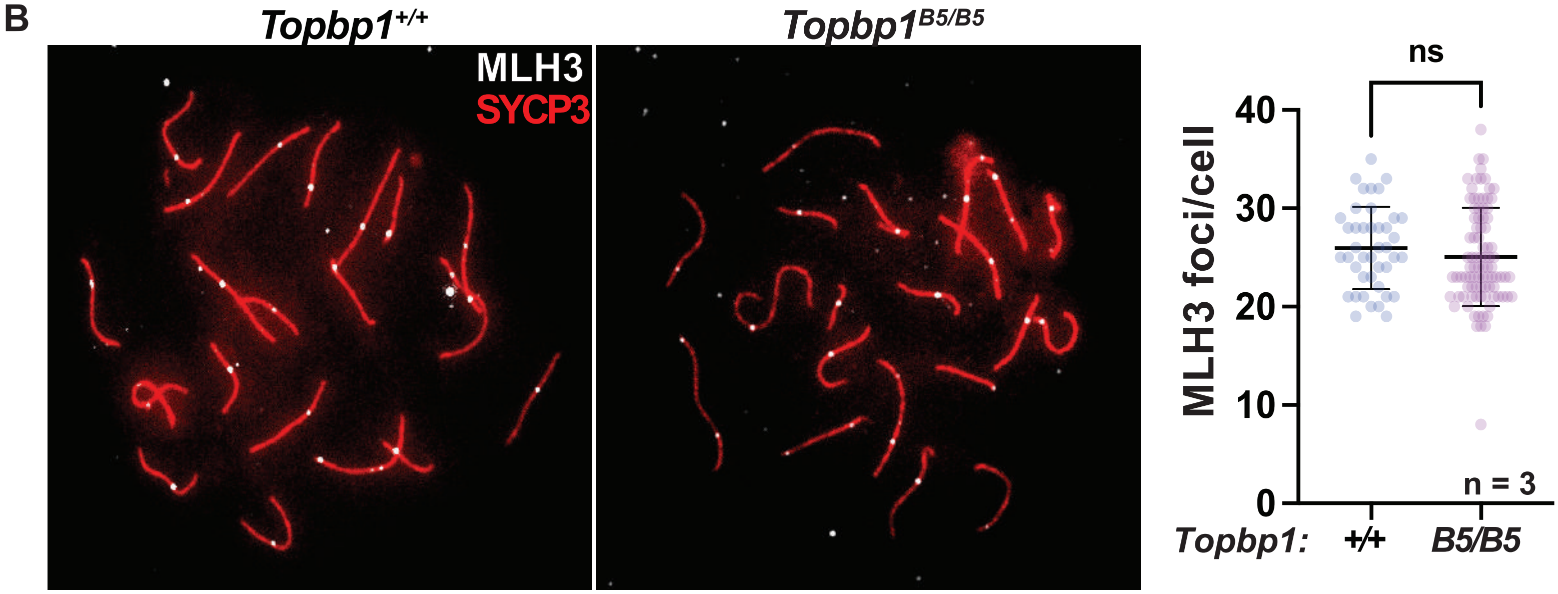

**Supplemental Figure 8**

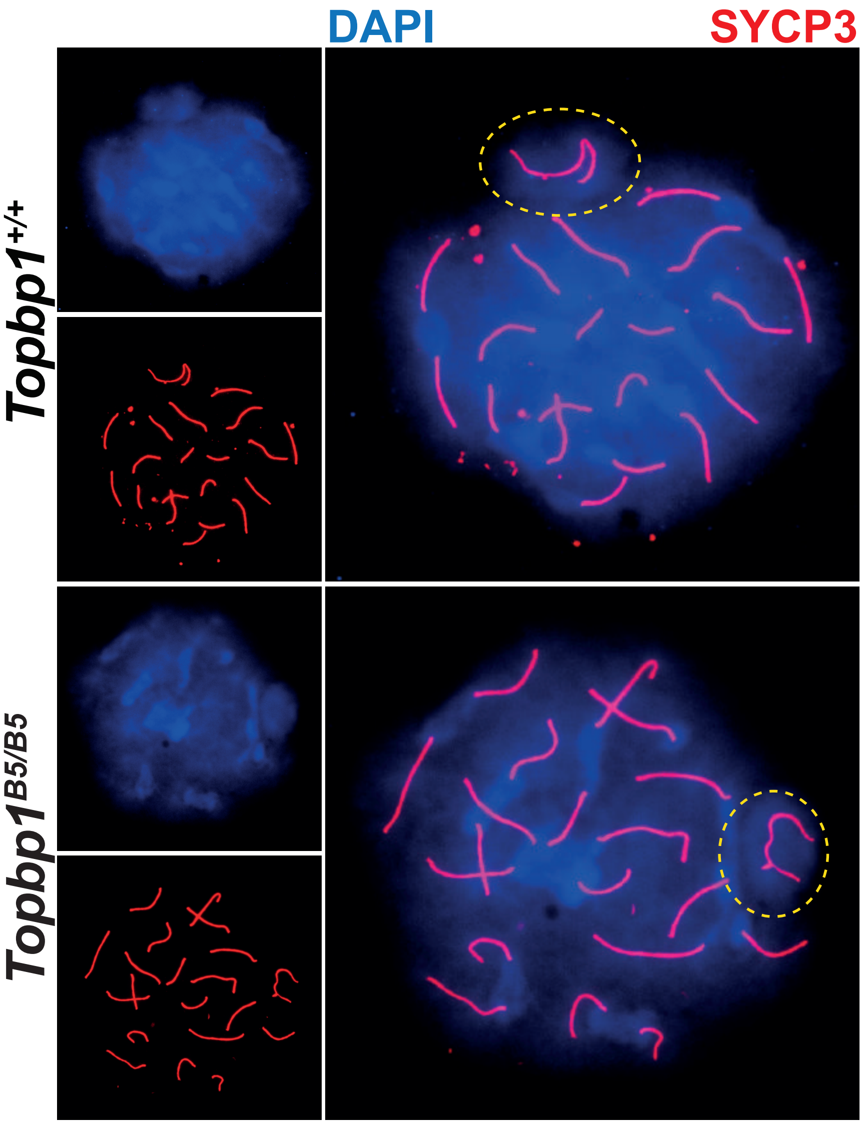

Supplemental Figure 9

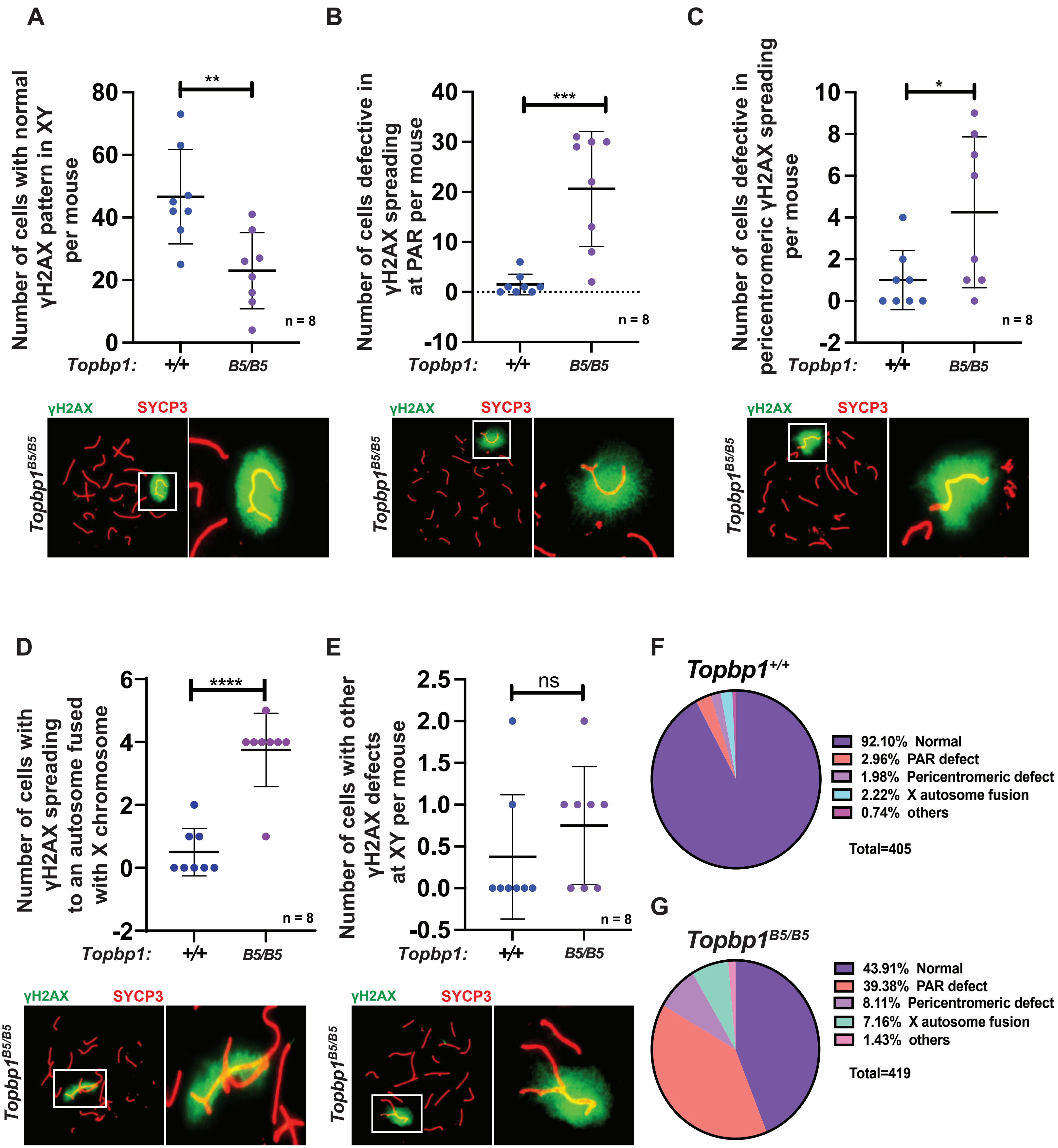

### Supplemental Figure 10

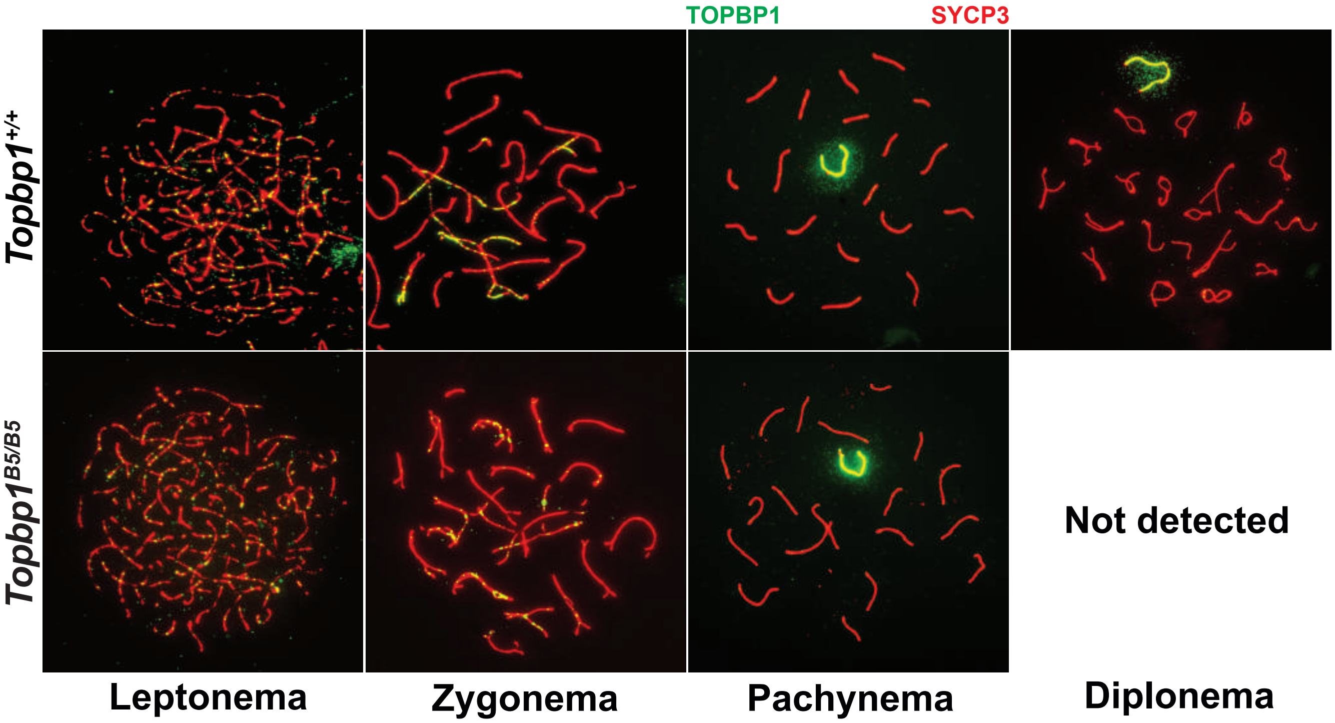

### Supplemental Figure 11

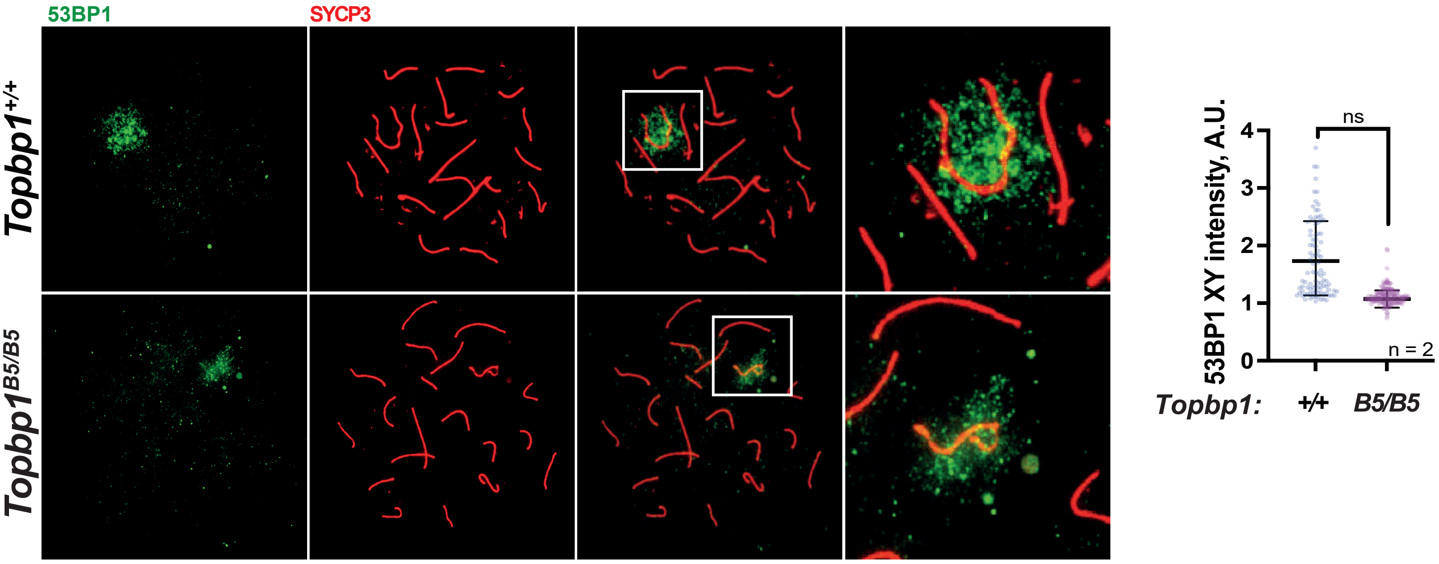

#### Supplemental Figure 12

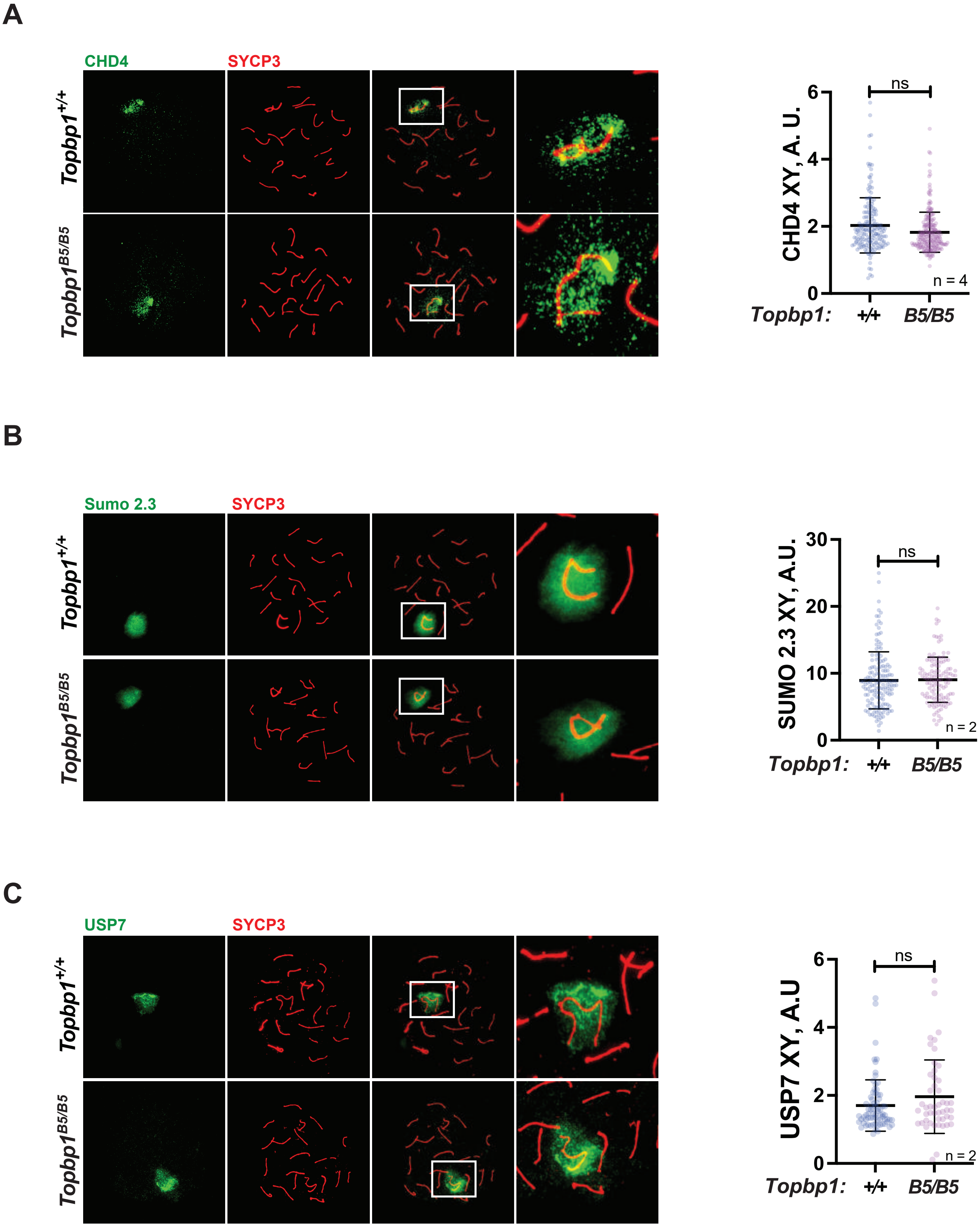

### Supplemental Figure 13

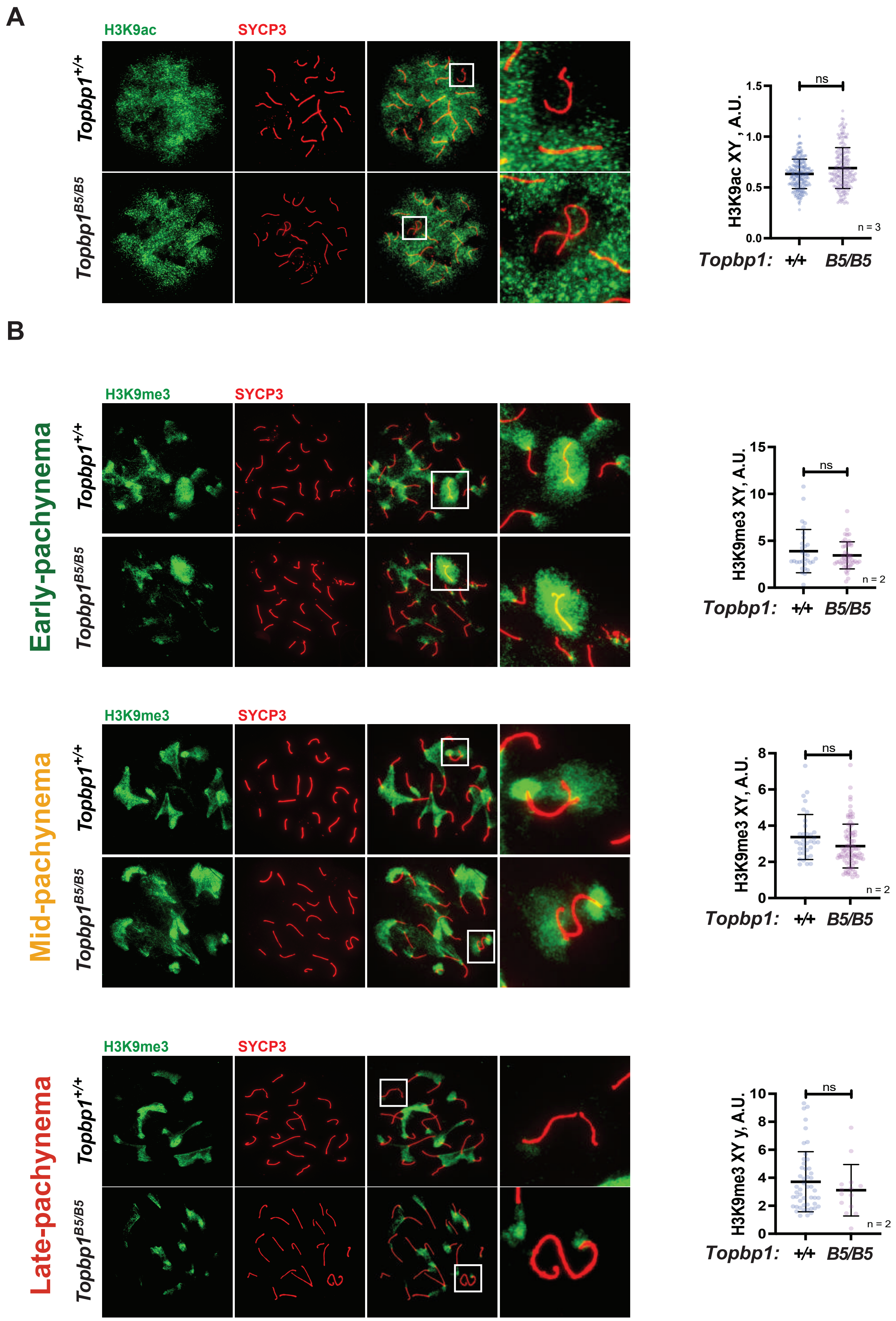

#### Supplemental Figure 14

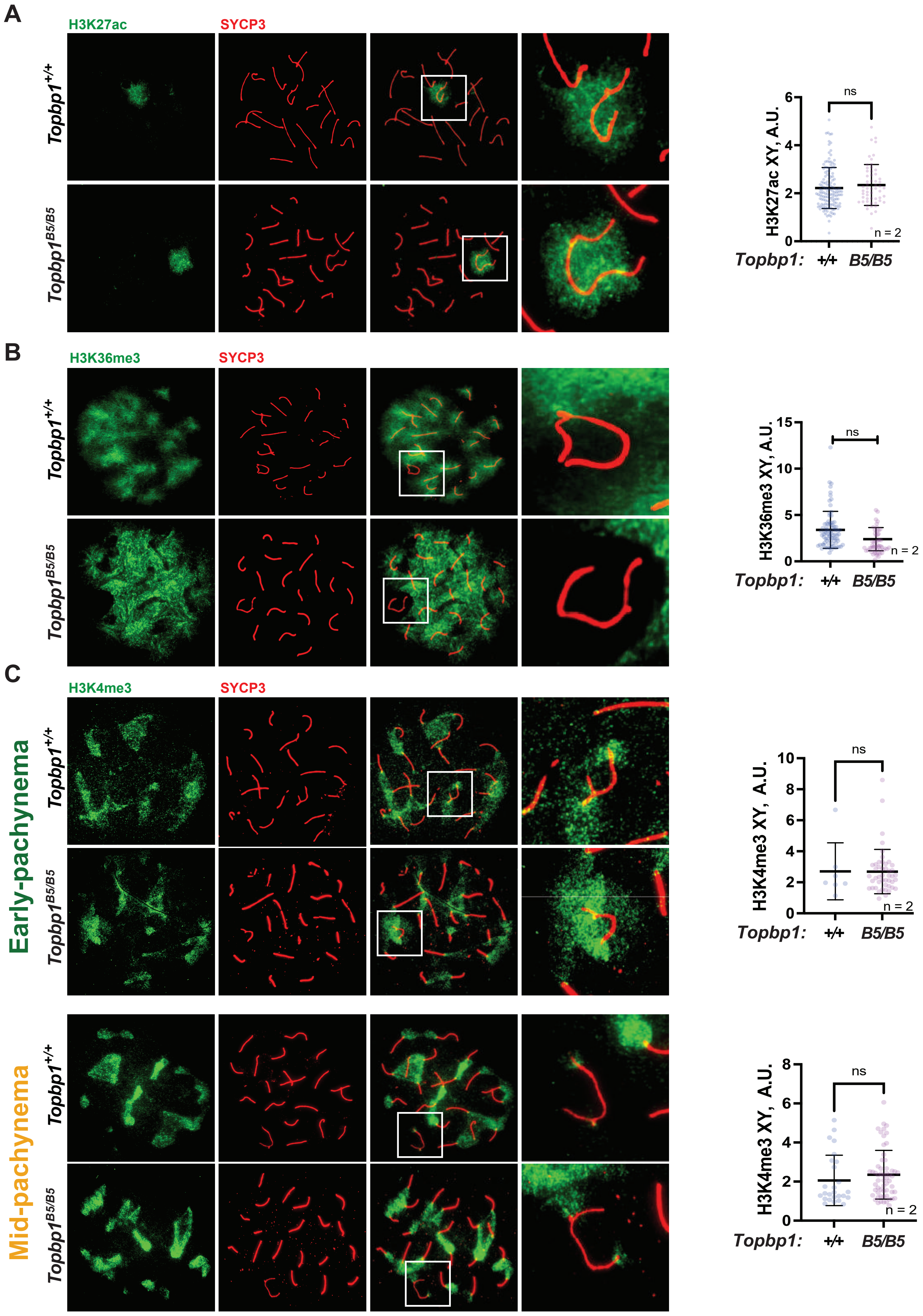

#### Supplemental Figure 15

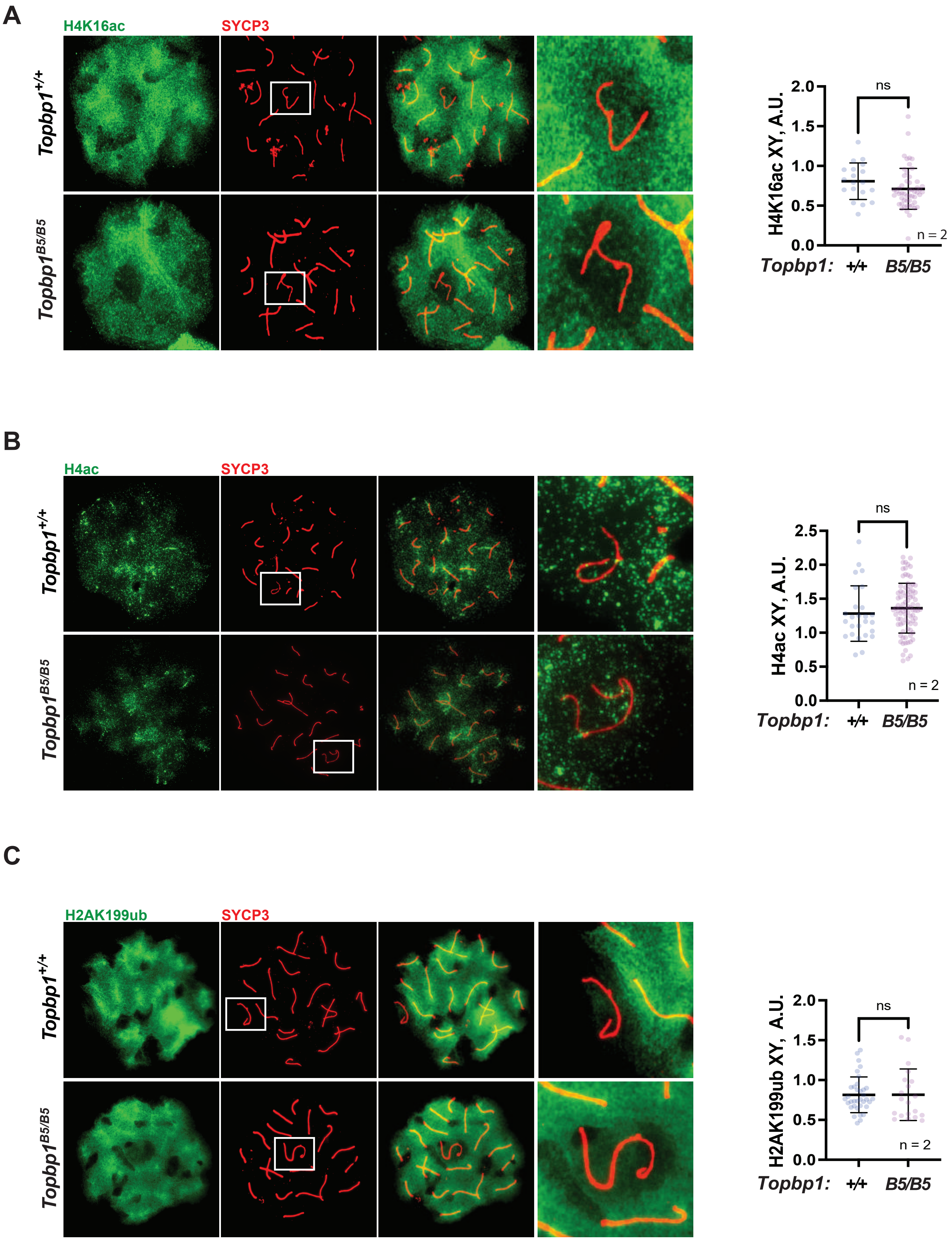

### Supplemental Figure 16

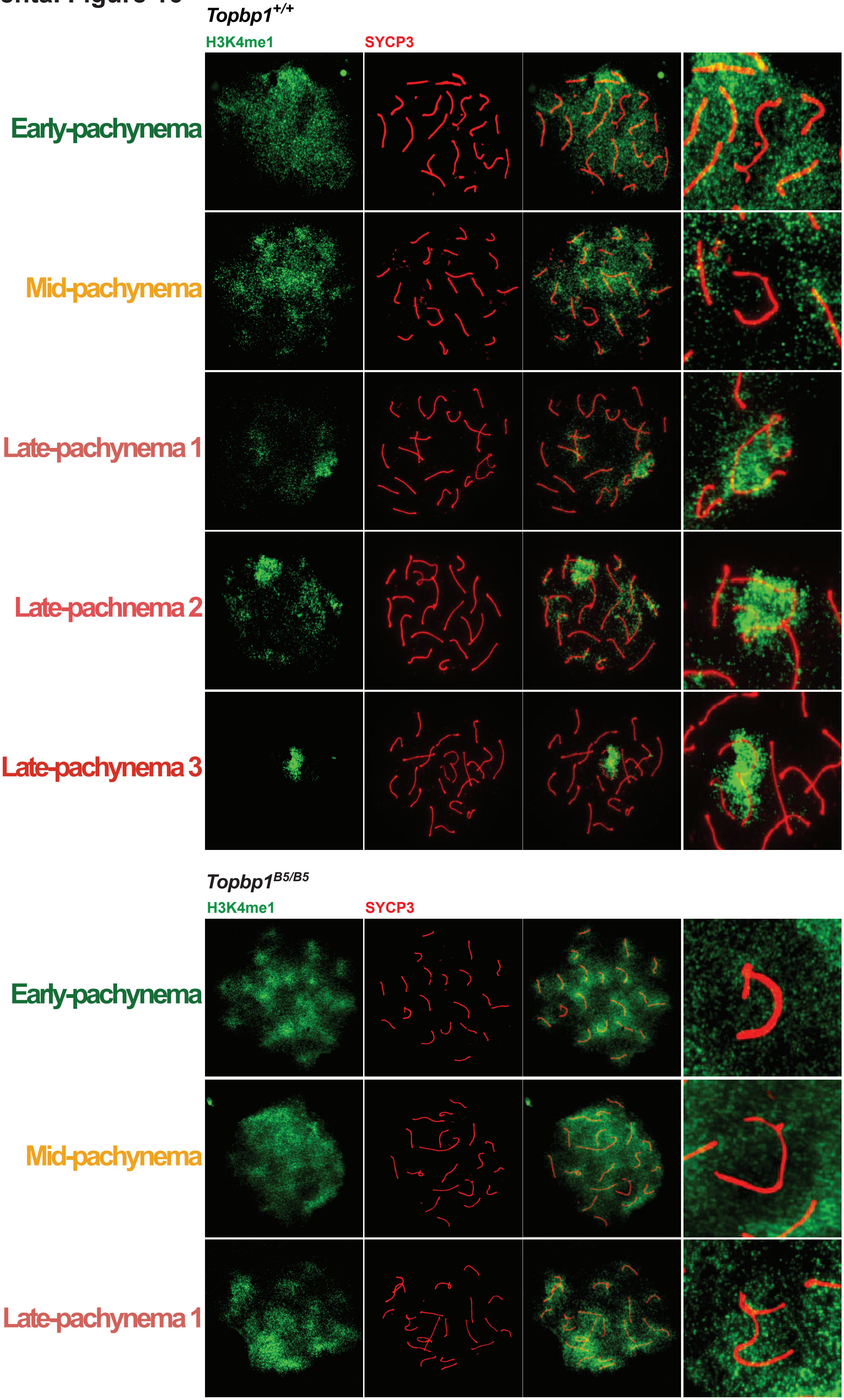

### Supplemental Figure 17

A

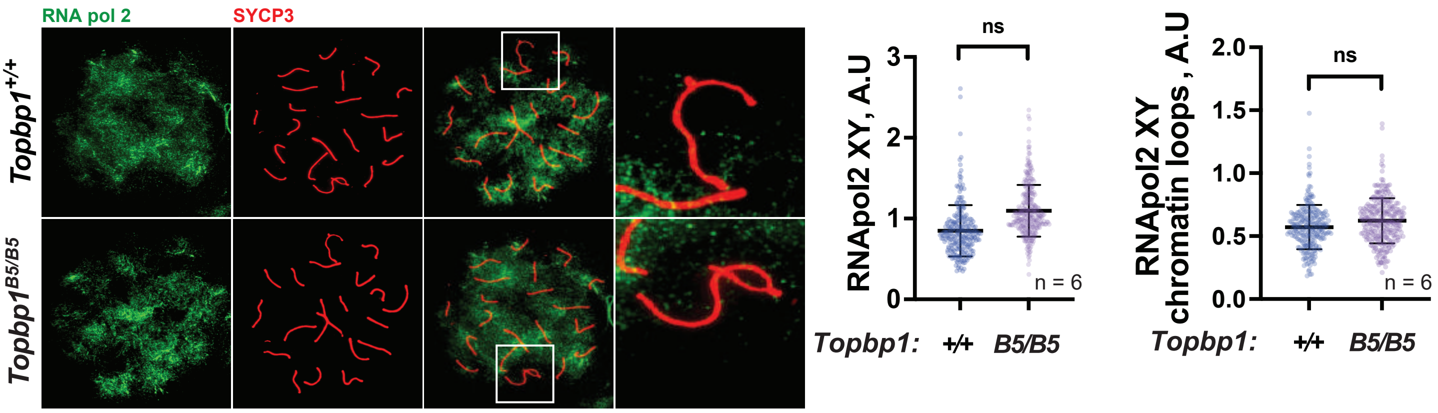


B

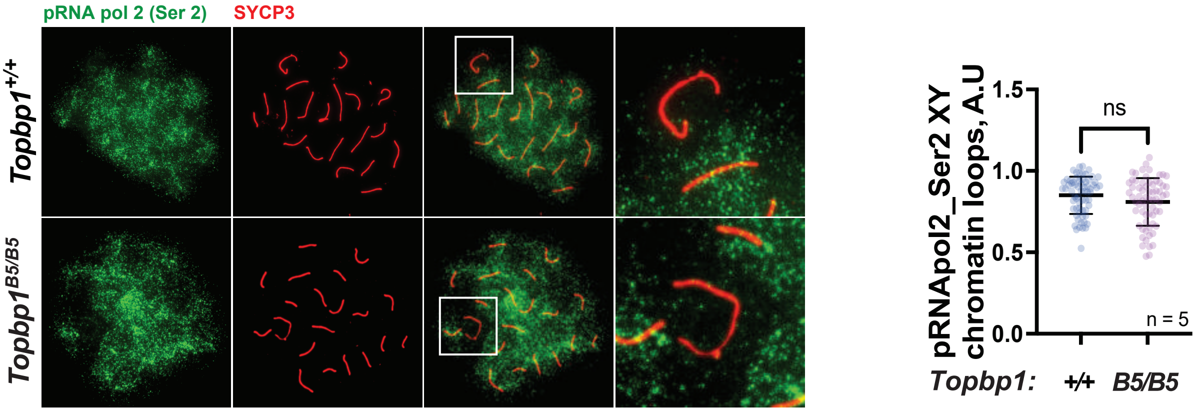


C

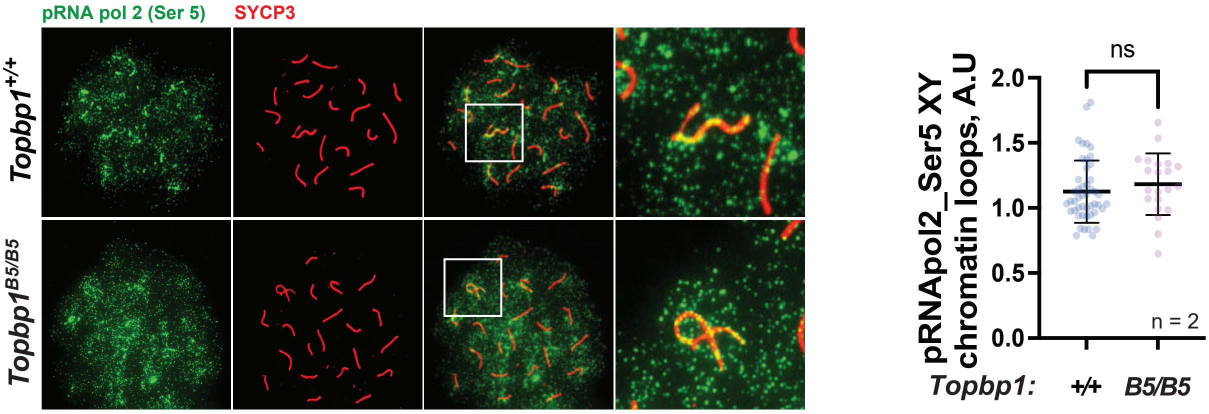

Supplemental Figure 18

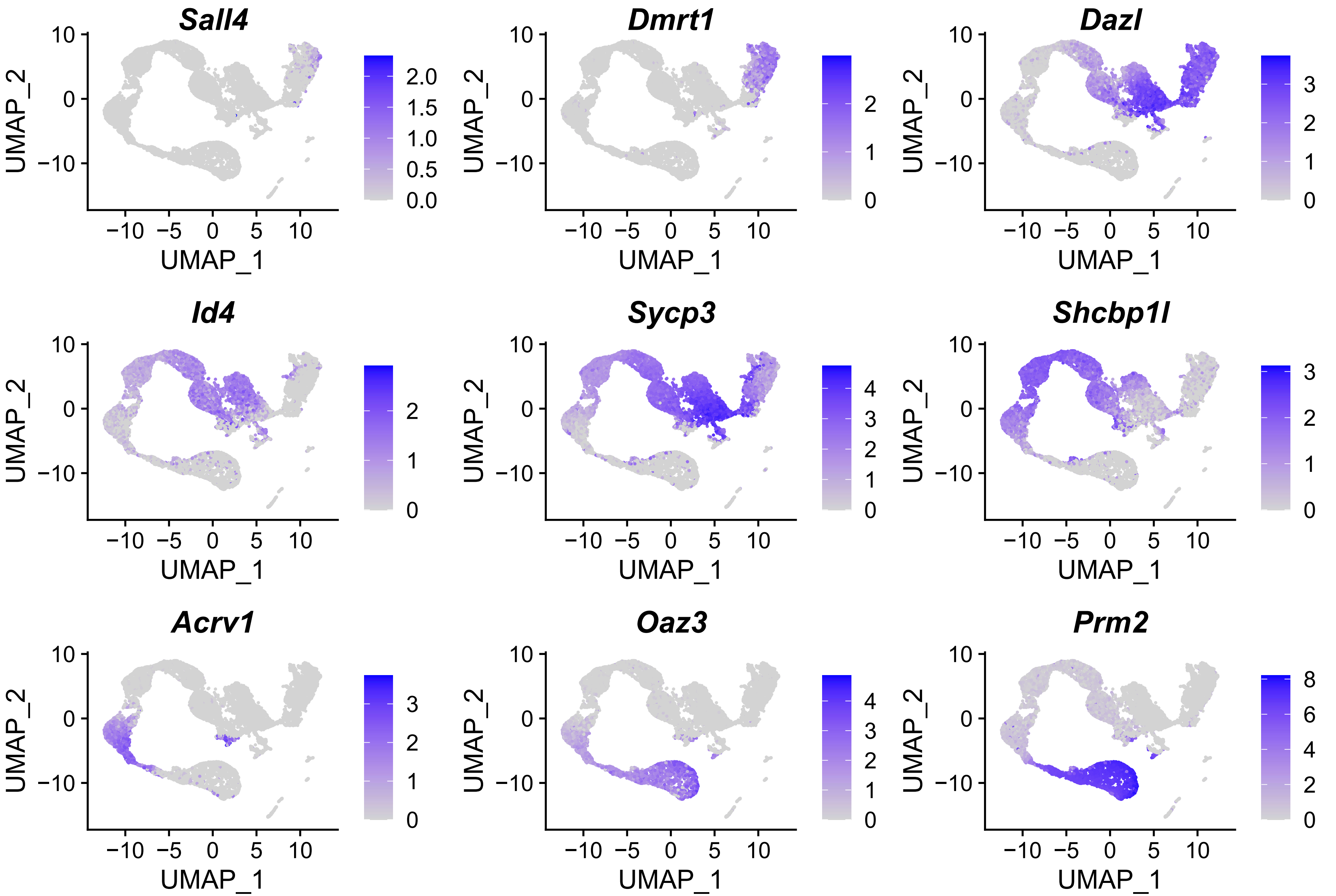

### Supplemental Figure 19

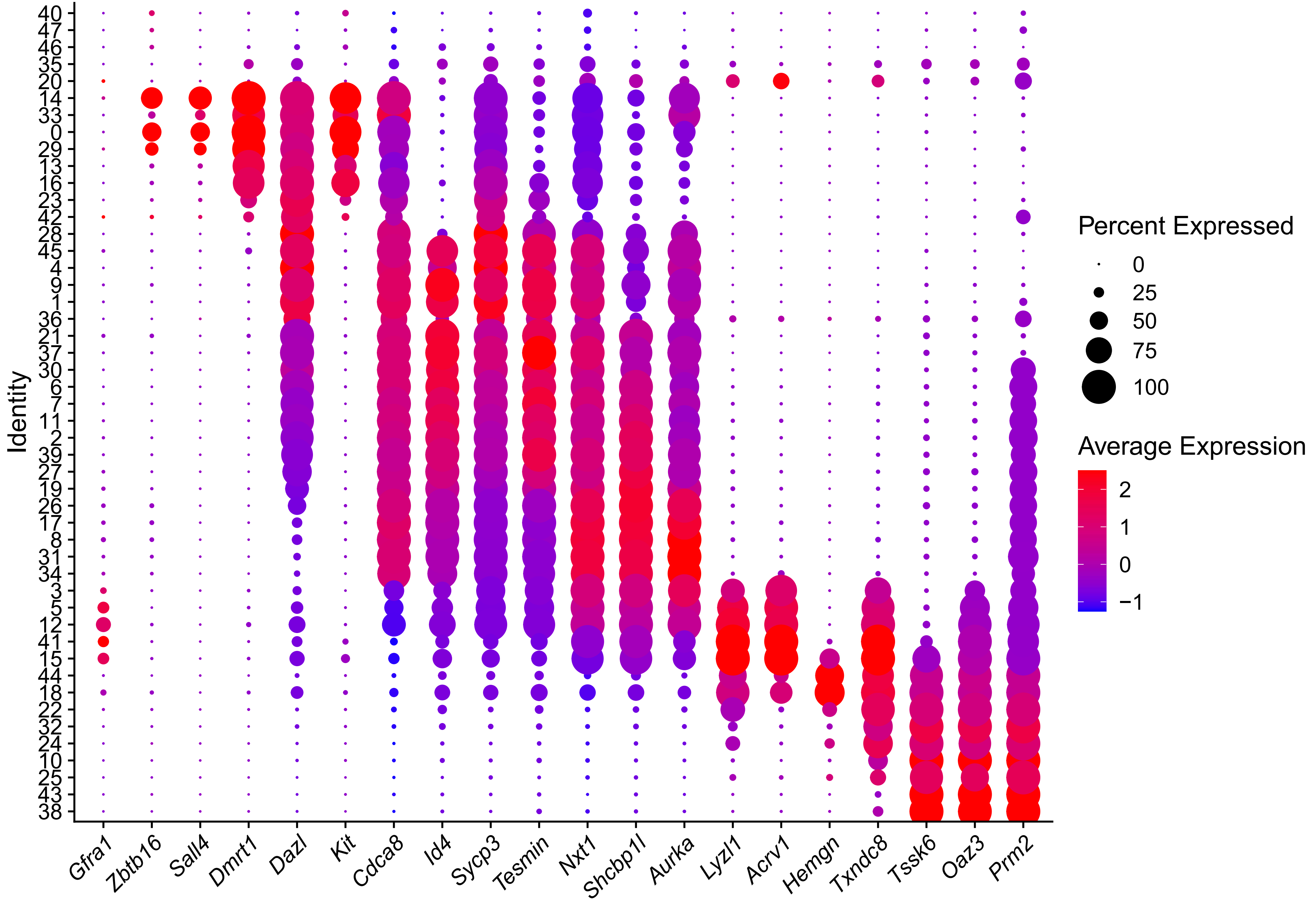

#### Supplemental Figure 20

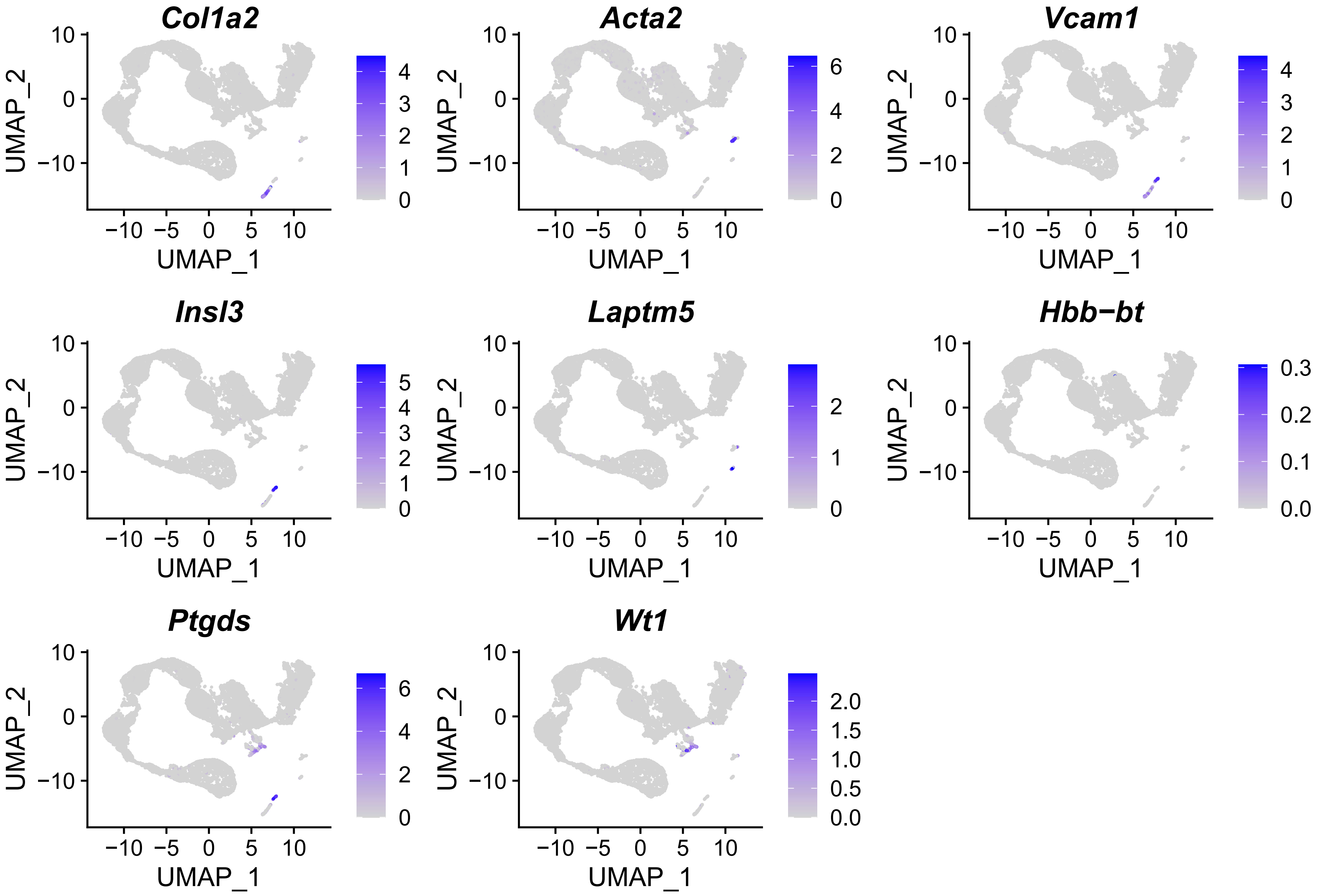

#### Supplemental Figure 21

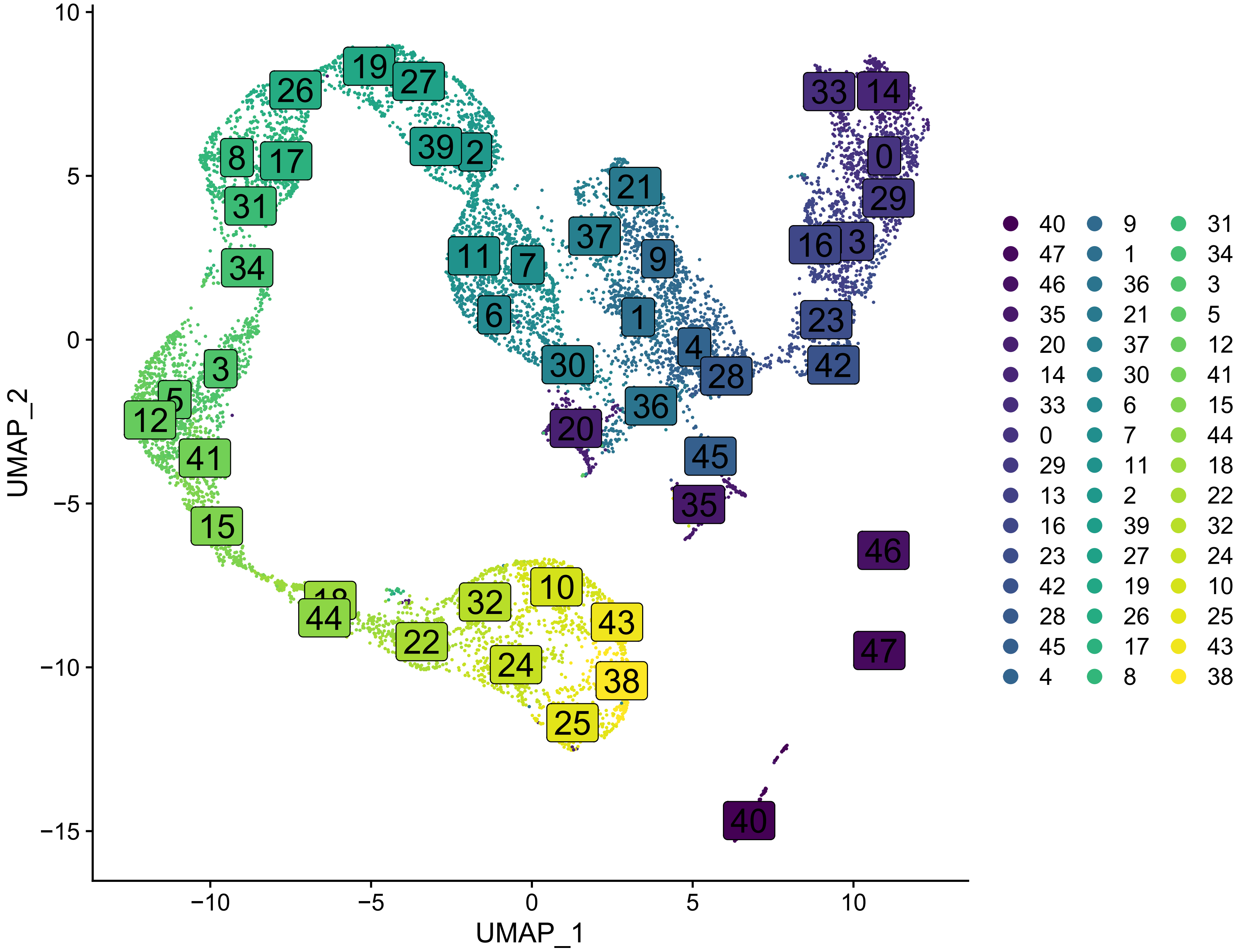

**Supplemental Figure 22**

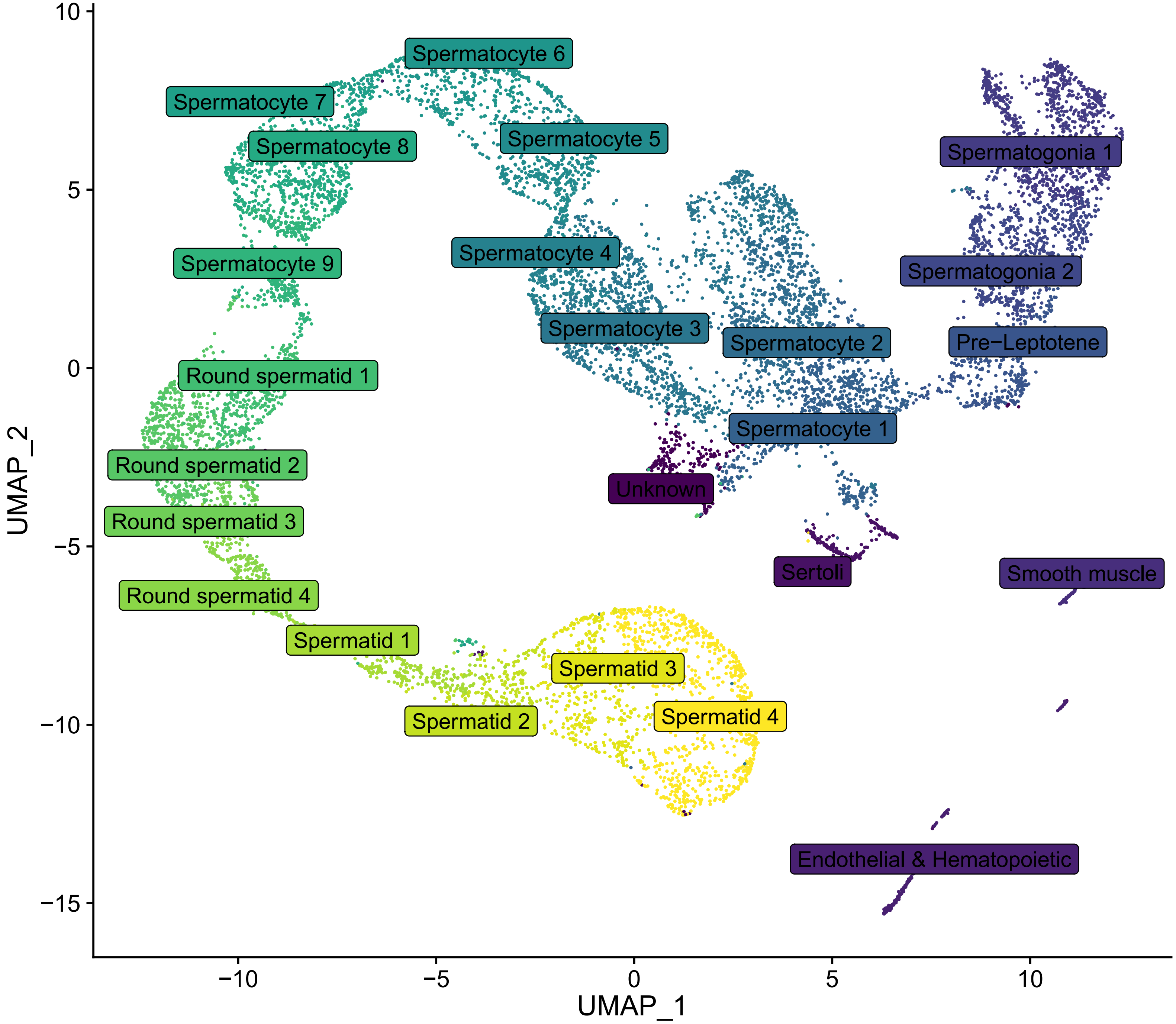

### Supplemental Figure 23

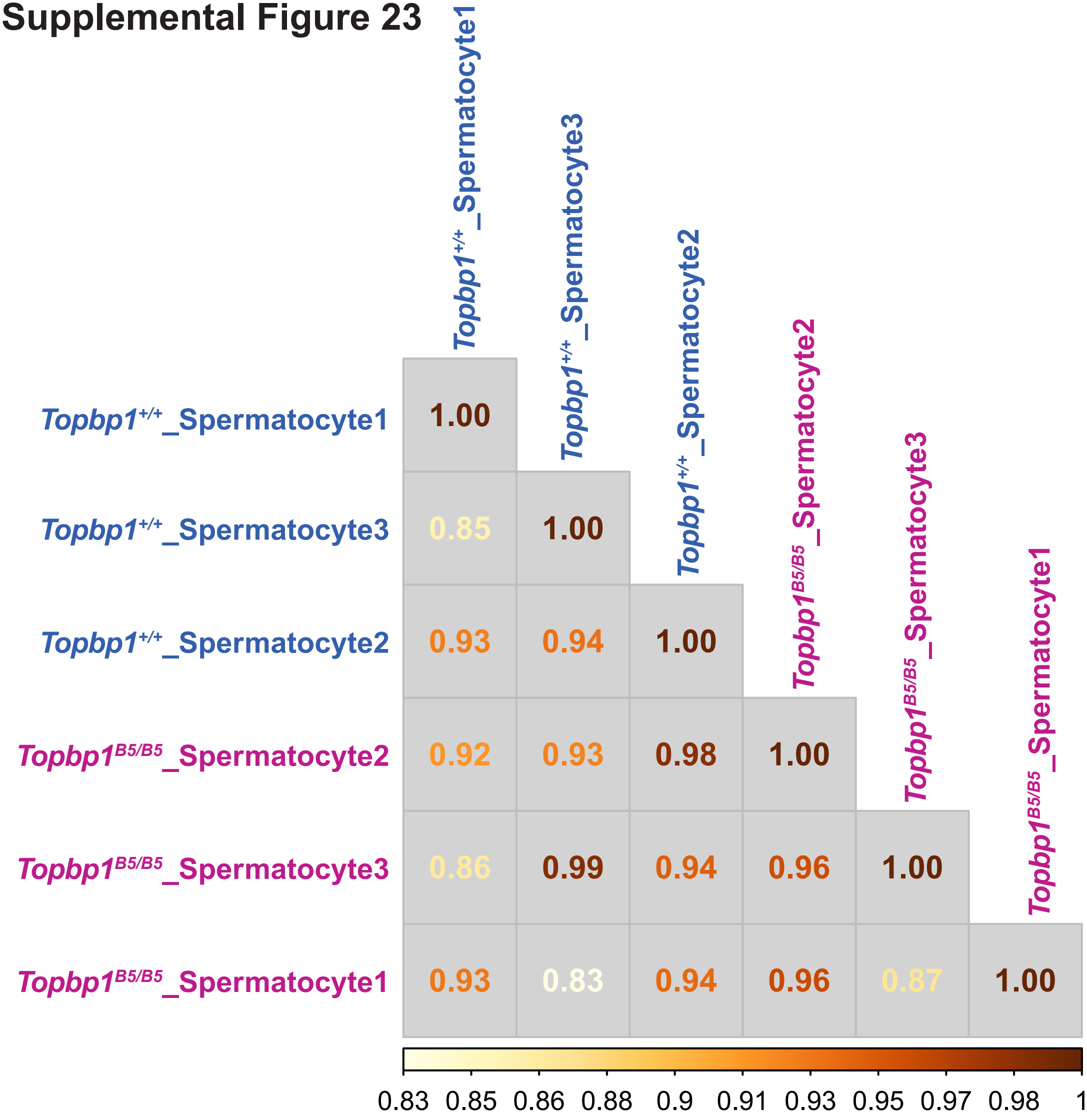

### Supplemental Figure 24

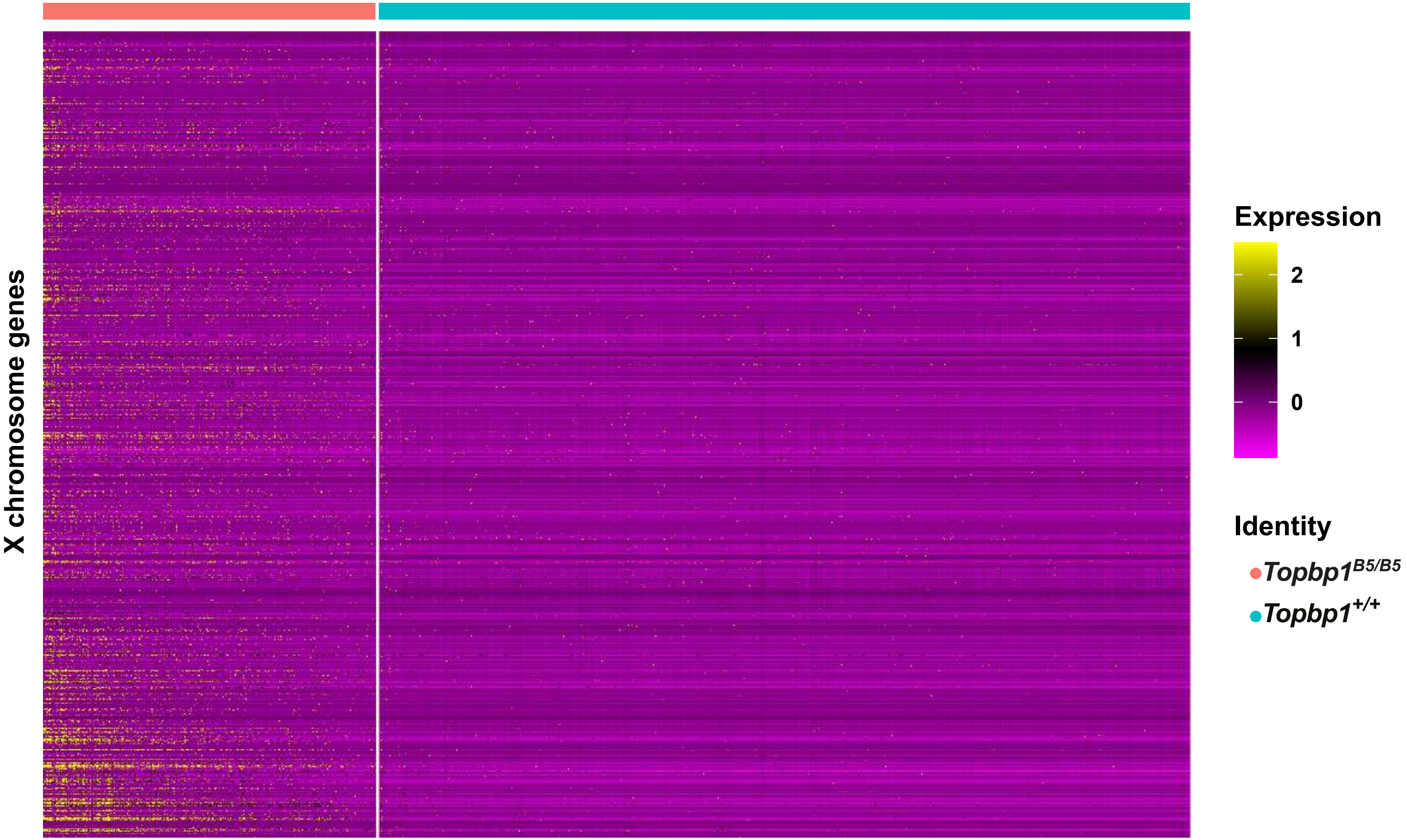

**Supplemental Figure 25**

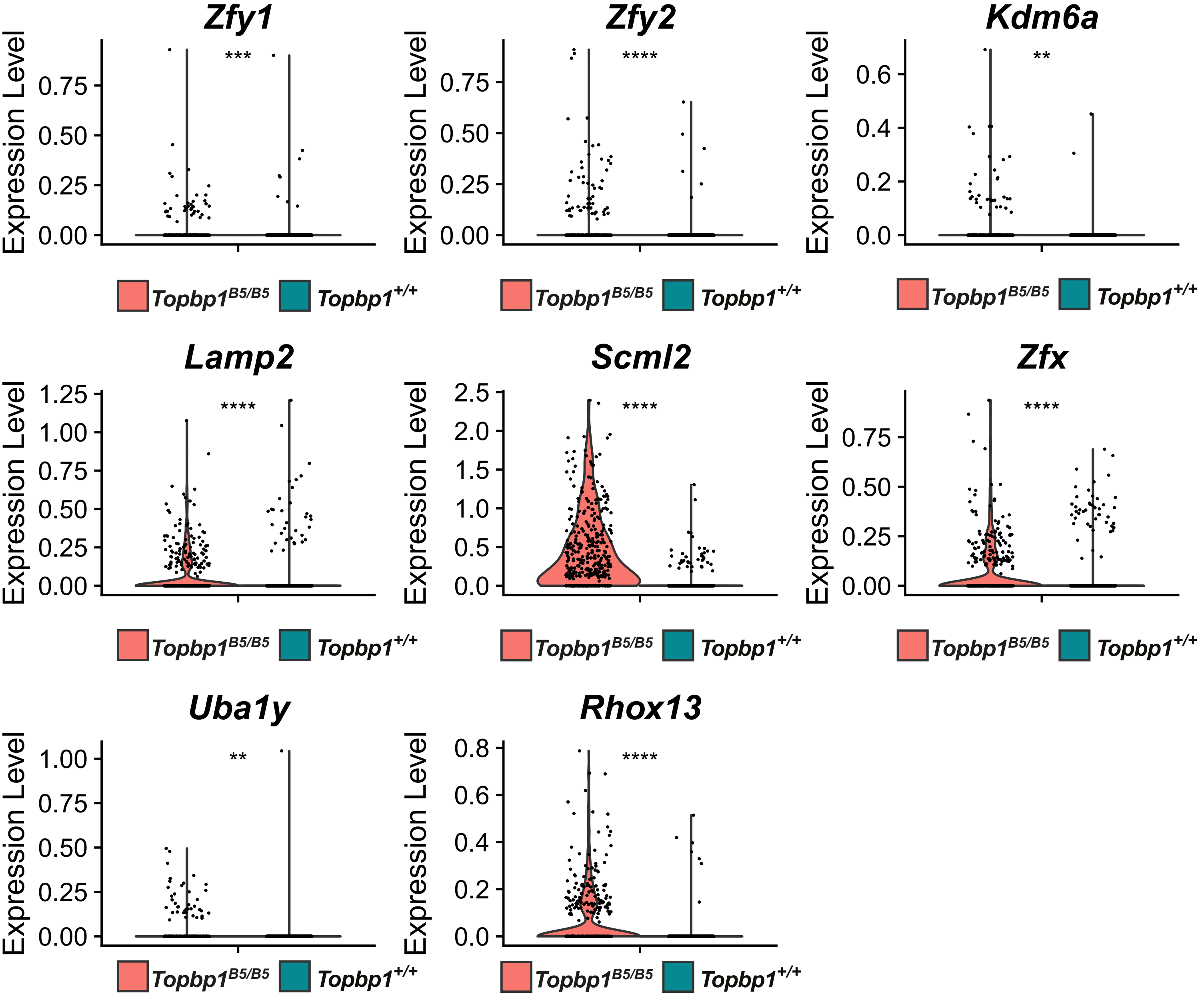

#### Supplemental Figure 26

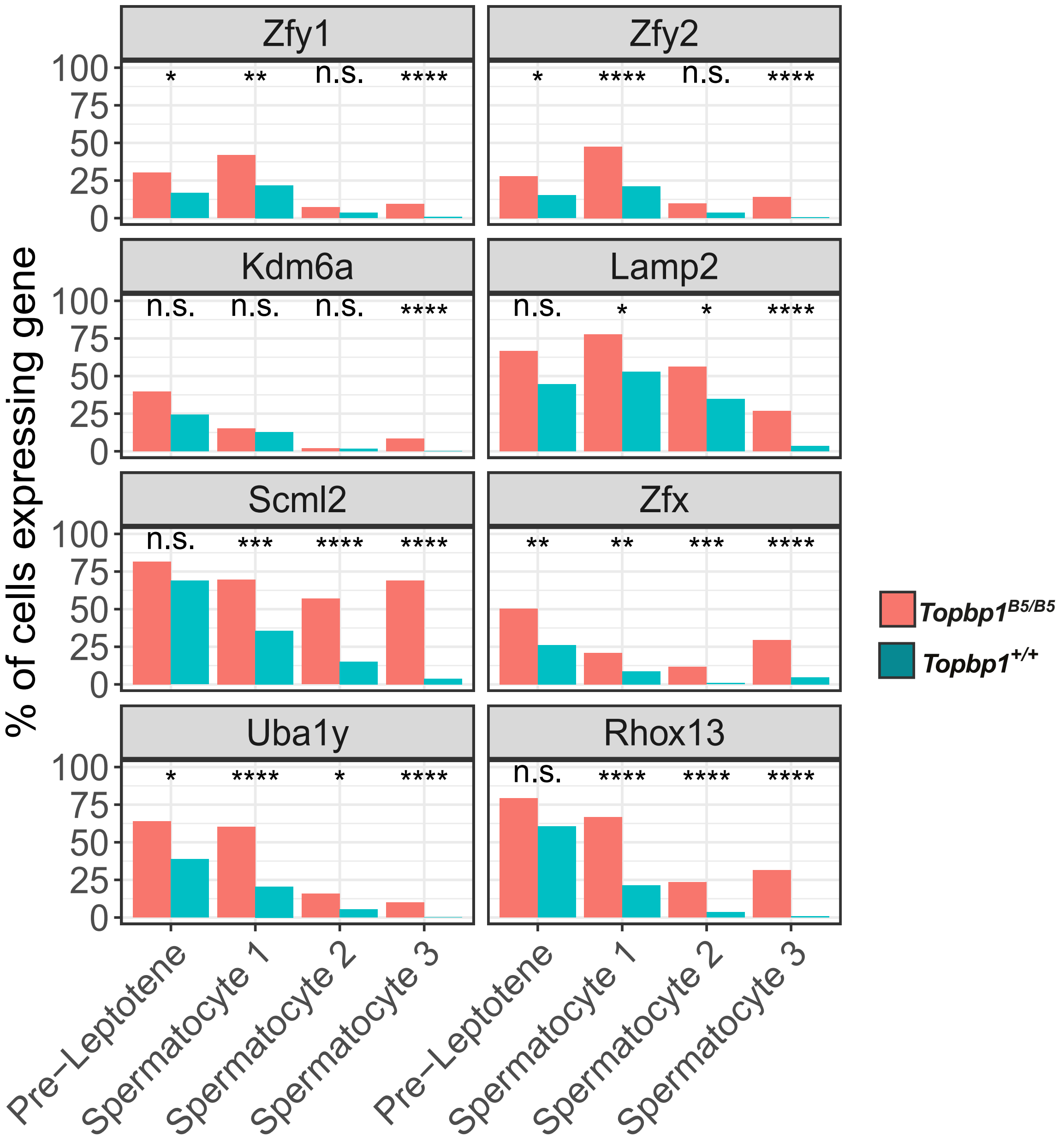

**Supplemental Figure 27**

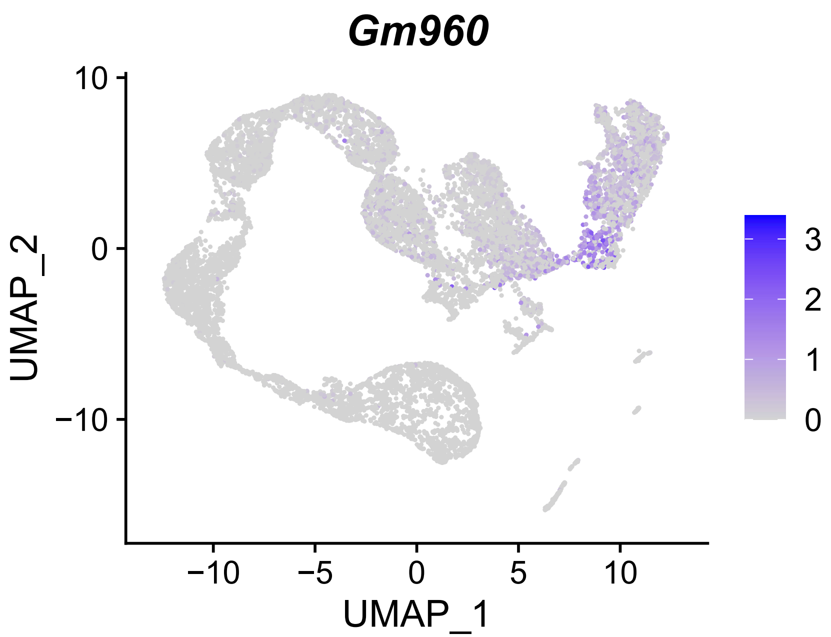

**Supplemental Figure 28**

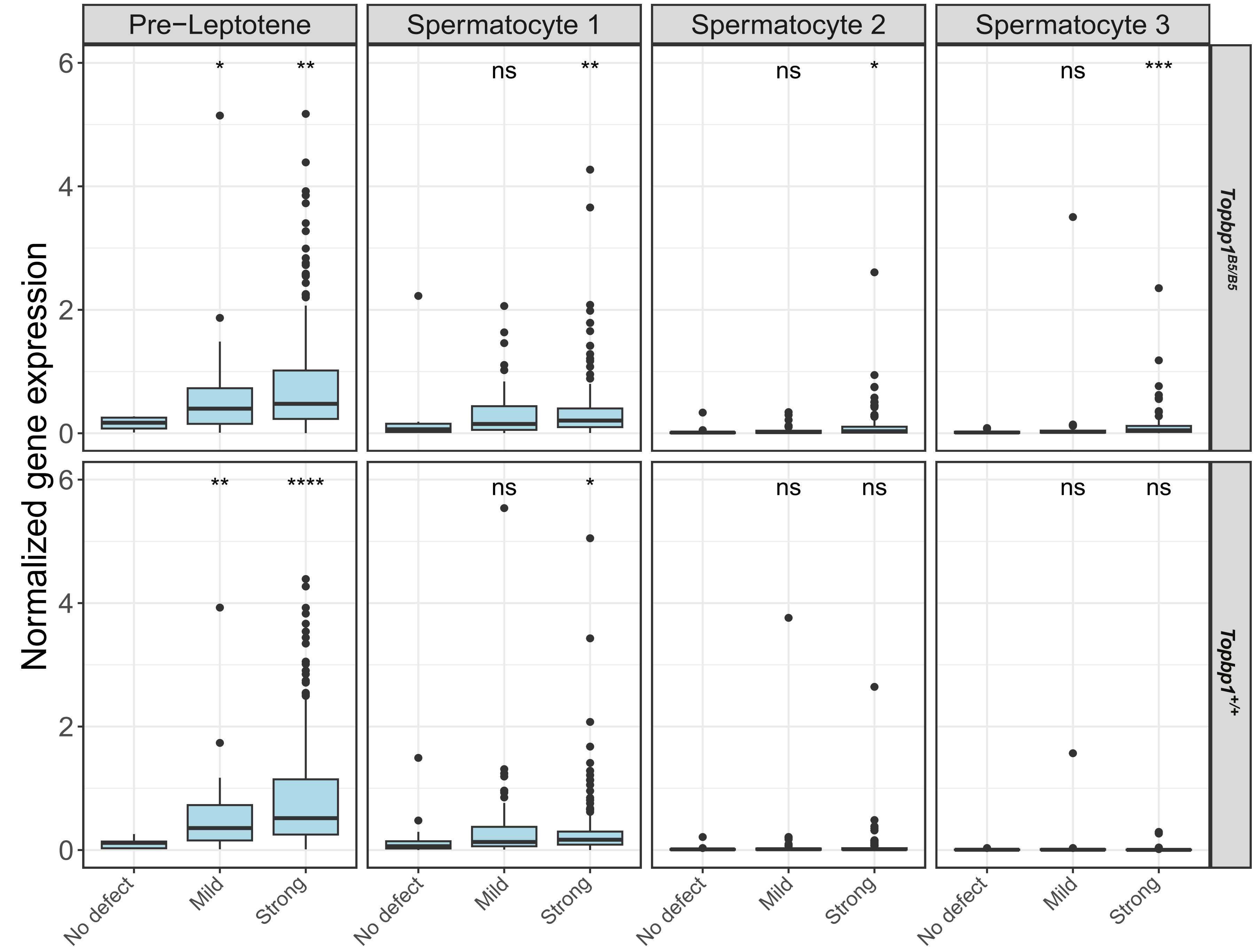

#### Supplemental Figure 29

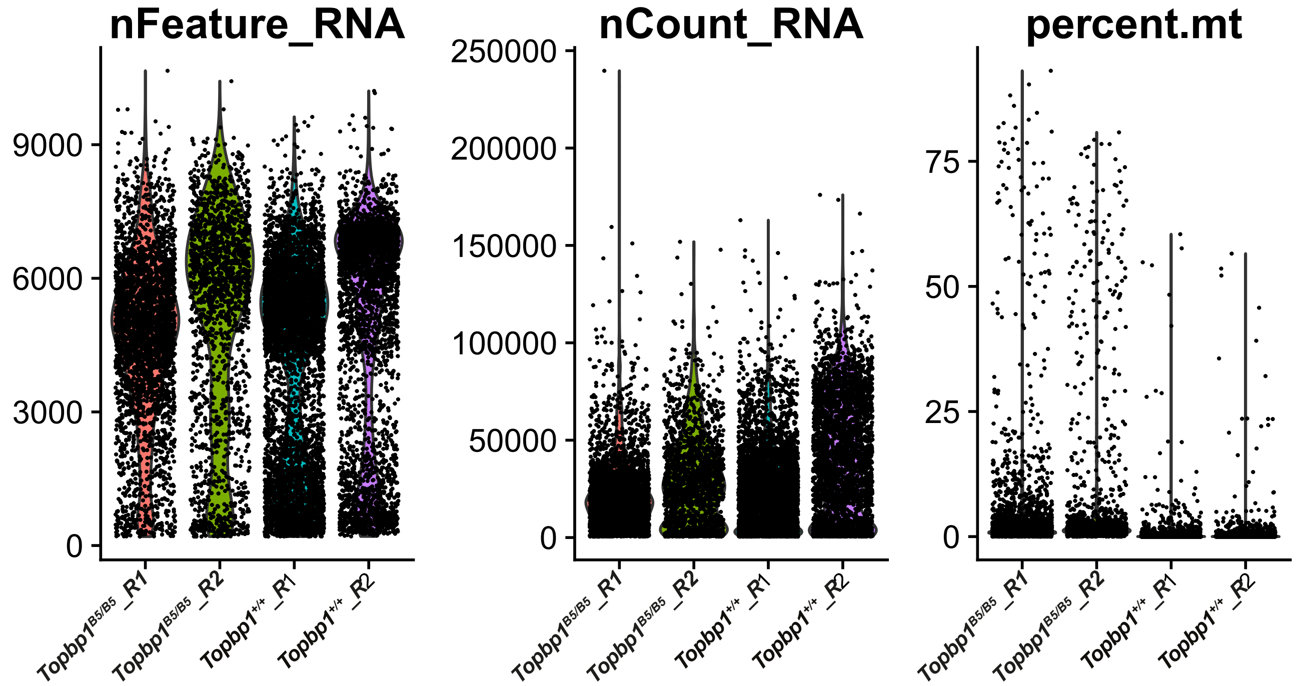
